# Heat-triggered remote control of CRISPR-dCas9 for tunable transcriptional modulation

**DOI:** 10.1101/606723

**Authors:** Lena Gamboa, Erick V. Phung, Haoxin Li, Jared P. Meyers, Gabriel A. Kwong

## Abstract

Emerging CRISPR technologies are enabling powerful new approaches to control mammalian cell functions, yet the lack of spatially-defined, noninvasive modalities to direct their function limit their potential as biological tools and pose a major challenge for clinical translation. Here we confer remote control of CRISPR-dCas9 activity using thermal gene switches, enabling the dynamic regulation of gene expression using short pulses of heat to modulate transcriptional commands.

## MAIN

RNA-guided endonucleases, which consist of Clustered Regularly Interspaced Short Palindromic Repeats (CRISPR) and CRISPR-associated proteins (Cas), have transformed genome engineering and are rapidly becoming indispensable tools in biomedical research. The programmable targeting capacity of Cas proteins has enabled applications that extend beyond genome editing, providing unprecedented new tools to control mammalian cell functions, including those which can regulate gene expression, modulate epigenetic landscapes, and manipulate chromatin structures^1^. These advances provide new opportunities for *in vivo* therapeutic applications, including recent demonstrations of dCas9 systems for gene therapy in rodent models of disease, such as diabetes, muscular dystrophy, and acute kidney disease^2–3^. Despite remarkable progress, significant hurdles remain that hinder practical applications of Cas technologies as *in vivo* tools and potential clinical therapies; these include off-target and off-tissue effects and the lack of precise methods to deliver or control Cas9 expression in target tissues. Leveraging CRISPR technologies to create platforms that noninvasively manipulate transcript levels and tune dosing based on short input signals may increase the effectiveness of targeted approaches for controlling synthetic cellular phenotypes *in vivo*.

Toward this end, several inducible systems have been developed to provide the ability to modulate the activity of Cas9 and its variants in living animals^4–5^. These include Cas9 systems that rely on chemical triggers using small molecule drugs, such as rapamycin and tamoxifen^6–8^, to activate and tune Cas9 activity by defined doses and at specific points in time. However, because these platforms rely on the systemic administration of chemical triggers to activate Cas-driven systems, they remain challenging to implement with spatial control. Alternatively, optical triggers offer noninvasive control of CRISPR-Cas9 that can be spatially defined by targeting with visible light^9–10^. Without invasive interventions, however, the efficacy of this approach is limited to superficial targets due to poor light penetration into biological tissue.

Here, we integrate heat as a remote trigger with the rapidly expanding CRISPR toolbox to confer tunable remote control of orthogonal transcriptional commands. In contrast to chemical or optical cues, pulses of heat can be delivered noninvasively with millimeter precision and at depth to anatomical sites by various approaches, such as infrared light^11^, high-intensity focused ultrasound^12^, or magnetic particles in alternating magnetic fields^13^. Recent control methods based on heat-induction to modulate gene expression^14–17^ include genetically encoded RNA thermometers^14^ and temperature-sensitive transcriptional regulators^16^. These systems have been developed to activate synthetic circuits in bacterial systems, but adaptation to mammalian cells have been limited due to practical concerns such as immunogenicity. Recently, we engineered mammalian thermal gene switches derived from the human heat shock protein HSP70B’ (HSPA6) locus to allow heat-triggered control of target gene expression^15^. In response to mild elevations in temperature (~40–42 ºC), our thermal gene switches undergo a sharp thermal transition to trigger transgene expression yet maintain negligible basal activity at body temperature^15^. Here we construct heat-sensitive dCas9 systems to provide the ability to reversibly and dynamically modulate mammalian transcription under remote thermal control.

To establish a thermal Cas9 transcriptional modulator, we cloned catalytically-inactive Cas9 (dCas9) variants (i.e. dCas9-VP64 and KRAB-dCas9) under the control of a thermal switch into lentiviral vectors (**Supplementary Figure 1**). dCas9 bears mutations in the RuvC1 and NHN nuclease domains^18^, rendering it unable to cleave DNA while retaining the ability to target specific sequences. When fused to protein domains, such as the Krüppel associated box (KRAB) or the tetrameric repeat of the herpes simplex viral protein 16 (VP64), dCas9 functions as a synthetic transcriptional regulator. In transduced and sorted HEK293T cells (Figure 1a; **Supplementary Figure 2**), we first sought to determine the ability to tune dCas9 expression levels by adjusting thermal trigger setpoints, such as temperature and heating duration. Upon mild hyperthermia, heat shock transcription factor 1 (HSF1), which is present as a monomer at basal temperatures, undergoes a conformational change that exposes hydrophobic interfaces to form trimers. HSF1 trimers then translocate to the nucleus to bind to heat shock response elements (HREs) to initiate transcription^15^. The formation of HSF1 trimers is affected by both the magnitude and duration of hyperthermic conditions, since changes in the conformational dynamics of monomeric HSF1 are temperature-dependent^19^. Therefore, to evaluate thermal responses, we measured KRAB-dCas9 expression 24 hours after heat treatment of 293T cells across distinct trigger temperatures (37– 42°C, Figure 1b) and heat pulse durations (0–60 min, Figure 1c). The ranges of these thermal conditions were selected to trigger HSF1 activation while maintaining cellular thermal tolerance and reversibility^15, 20^. As anticipated, increasing the trigger temperature to values above 37°C resulted in a sharp thermal transition that significantly increased dCas9 expression at temperatures greater than 41°C (\*\*\*\**p* < 0.0001) by greater than 7-fold (Figure 1b, d), consistent with thermal switch control of reporter genes previously reported^15^. We observed a similar increase in thermal response as the heating durations were progressively extended from 0 to 60 min while maintaining a constant trigger temperature of 42°C (Figure 1b). Compared to optogenetic CRISPR-Cas9 tools that require continuous light illumination to maintain “ON” state activity^9–10^, we observed that heat activation of our thermal switch by as little as 15 minutes significantly elevated and maintained dCas9 expression (Figure 1b,d \*\*\*\**p* < 0.0001).

**Figure 1.**
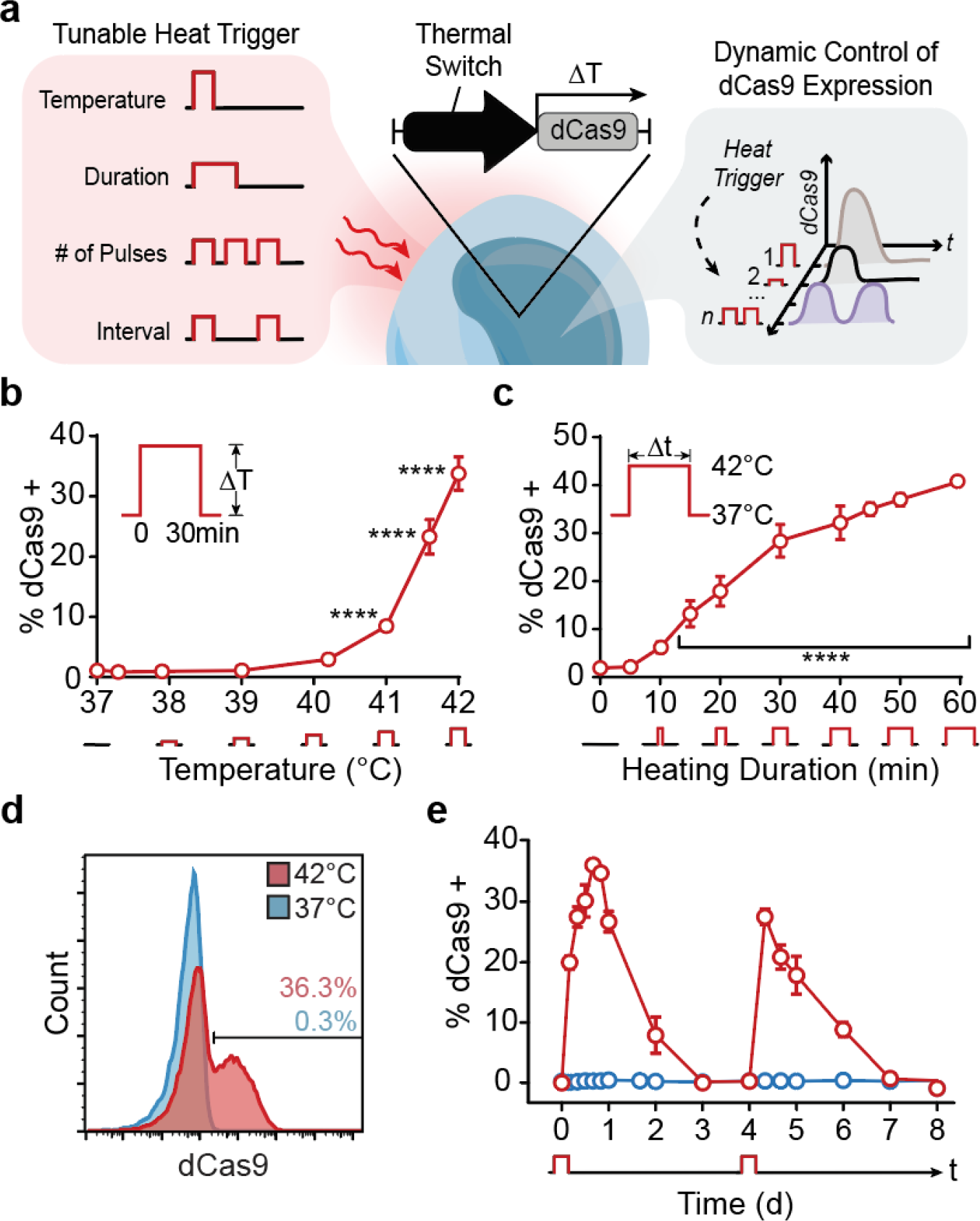
Heat-triggered gene switch enables dynamic, tunable control of dCas9 expression. (a) Following exposure to distinct thermal triggers, which are tunable by the modulation of temperature, heating duration, the number of pulses delivered, and time interval between heating doses, cells transduced with a thermal switch dynamically express dCas9 variants. KRAB dCas9 expression after (b) increasing activating temperature from 37°C to 42°C or (c) heating duration from 0 to 60 min (*n=*3, mean ± s.d. one-way ANOVA, \*\*\*\**p*<0.0001). (d) KRAB-dCas9 expression in stably transduced HEK293T cells following 30 min of heating (*t* = 16 hr) (c) Kinetic trace of KRAB-dCas9 expression following 30 min of heating at 42°C treated at t = 0 d and t = 4 d (*n=*3, mean ± s.d., blue trace=37°C, red trace = 42°C).

To evaluate longitudinal control, we monitored dCas9 expression following 30-minute pulses of heat (42°C) at *t* = 0 d and *t* = 4 d. In contrast to unheated controls which showed negligible levels of dCas9 (<0.43%), we detected significant increases in dCas9 expression over the course of 3 days after each heating cycle with similar expression and decay half-lives (Figure 1e, t_1/2_ = ~20 hrs). Similar switch-on and switch-off kinetics were observed by thermal control of catalytically-active Cas9 (**Supplementary Figure 3**). Collectively, these data demonstrate that the rapid and transient expression of Cas proteins can be sustained for several days with the delivery of discrete, short pulses of heat minutes in duration.

Having demonstrated the use of heat to trigger expression of Cas9 constructs, we next explored thermal modulation of transcriptional activity. To control gene activation, we co-delivered plasmids encoding for the thermal switch, the MS2:P65:HSF1 (MPH) activation complex, and multiple sgRNAs targeting either the human *IL1RN*, *GzmB*, or *CCL21* promoter (Figure 2a, **Supplementary Figure 1a**, **Supplementary Table 1**). We included MPH due to its ability to increase activation efficiency when it binds to MS2 loops on the sgRNA backbone^21^. *IL1RN*, *GzmB*, and *CCL21* were chosen for their diversity in function, as well as for their important roles across biological processes, such as cell signaling, apoptosis, and cell migration^22–24^. In accordance to our results that increasing trigger temperature leads to higher levels of dCas9 protein, elevated temperatures enhanced *IL1RN*, *GzmB*, and *CCL21* activation by up to 4-fold 72 hrs after heating (Figure 2b, **Supplementary Figure 4**). Notably, hyperthermia alone controls did not induce upregulation of *IL1RN*, *GzmB*, or *CCL21*, demonstrating that transcriptional upregulation reported is due to thermal control of dCas9 and not endogenous cell responses to heat (**Supplementary Figure 5**).

**Figure 2.**
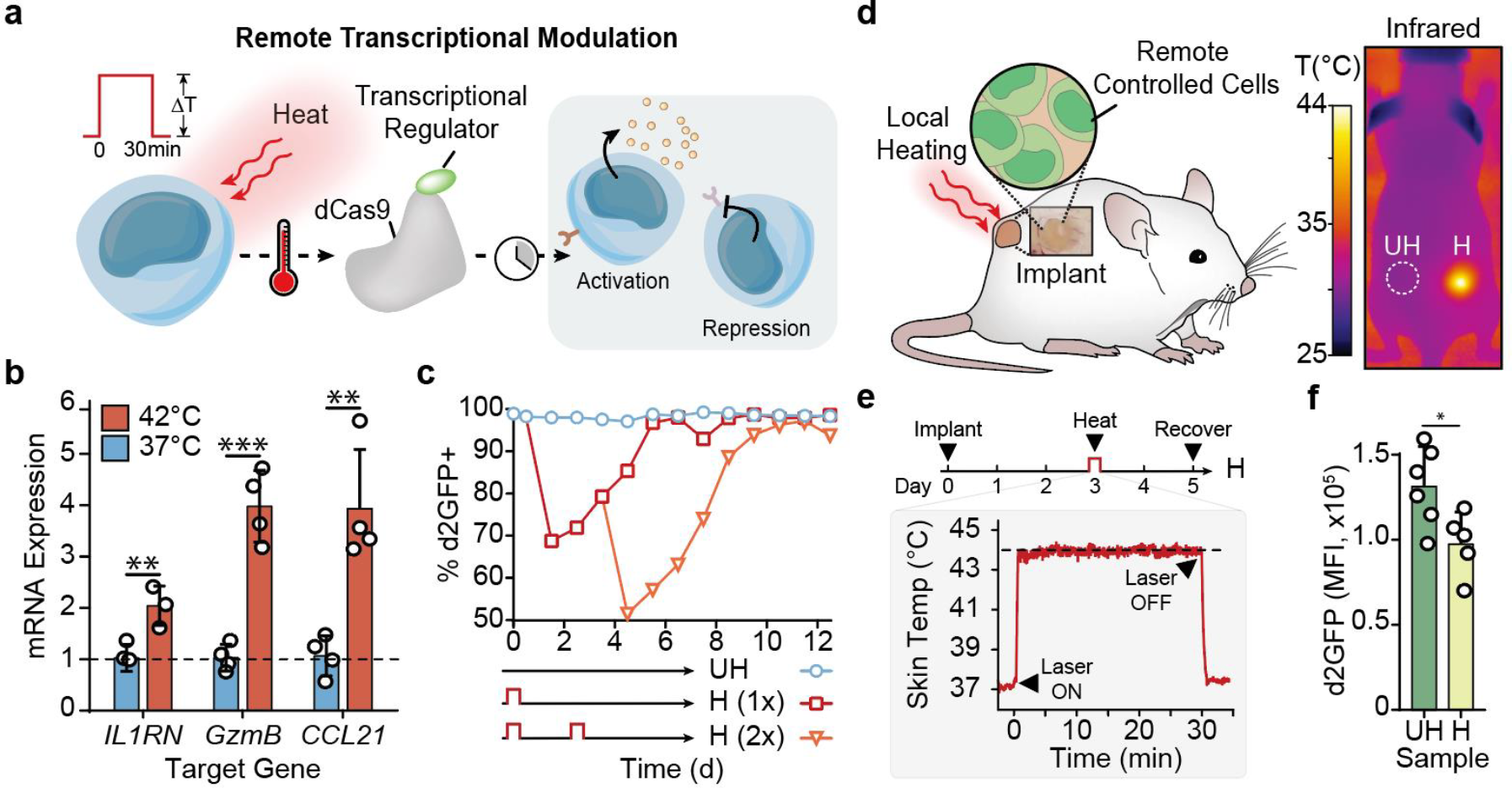
Modulation of mammalian cell transcription by remote-controlled thermal switches. (a) Thermal switches in HEK293T cells are triggered by a 30 min heat pulse, enabling the remote modulation of target gene transcription. (b) Endogenous gene activation as assessed by qRT-PCR 72 hrs after heating at the 42°C (*n* = 3-4, mean ± s.d., unpaired t-test, \*\**p*<0.01, \*\*\**p*<0.001). (c) Kinetic trace of d2GFP suppression in HEK293T cells following 0, 1, or 2 heating doses (*n* = 3, error bars show s.d. and are smaller than the displayed data points). (d) Left: Engineered cells were embedded in Matrigel tissue phantoms, subcutaneously injected in the rear flank of mice, and subsequently heated locally using near-infrared laser light. *Right*: Thermal images of a mouse undergoing laser-induced plasmonic heating (H = Heated, UH = Unheated). (e) Representative kinetic thermal trace showing skin temperature of 3 × 3-pixel ROI centered on implant site. Triangle indicates timepoint when laser is turned on or off. (f) Mean fluorescence intensity of d2GFP in HEK293T cells recovered from 37°C and 42°C heated implants (*n*=5-6, mean ± s.d., unpaired t-test, \**p*<0.05).

To direct transcriptional suppression with dCas9 (**Supplementary Figure 1b**), we targeted destabilized eGFP (d2GFP) with a 2-hour half-life to assess the kinetics of remote-controlled gene suppression to preclude confounding signals from long-lived reporters. In order to select the most potent d2GFP-targeting sgRNA sequence, we screened four sgRNA candidates using a catalytically-active Cas9 and found that the top performing guide (A3) knocked out d2GFP in greater than 70% of the cell population (**Supplementary Figure 6**, **Supplementary Table 2**). To test thermal control of transcriptional suppression, cells containing the thermal switch driving KRAB-dCas9 expression were transduced with d2GFP and the A3 d2GFP-targeting sgRNA sequence. In the absence of a thermal trigger, cellular expression of d2GFP was constant and maintained for over 10 days with negligible suppression detected (>97% of cells remained d2GFP+). By contrast, we observed that heated cells significantly suppressed d2GFP (Figure 2c, **Supplementary Figure 8a**) and the level of suppression could be directly modulated using multiple doses of heat or treatment windows – for example, delivery of 30-min pulses administered 3.5 days apart resulted in further suppression of d2GFP expression (**Supplementary Figure 8b-d**). Under all conditions, gene suppression was reversible and full recovery of reporter expression was observed within several days after the last heat treatment.

After demonstrating transcriptional modulation *in vitro*, we next set out to implement this system for remote thermal control of gene expression *in vivo*. Spatially controlled delivery of heat is routinely employed in thermal medicine to improve treatment efficacy of drugs and can be accomplished by various modalities such as plasmonic photothermal heating^11–13^. To implement this system *in vivo*, we subcutaneously implanted tissue phantoms in the rear flank of nude mice (Figure 2d). These tissue phantoms comprised Matrigel implants seeded with plasmonic gold nanorods (AuNRs), which absorb near infrared (NIR) light and convert the resonant energy to heat^11^, and d2GFP+ HEK293T cells containing d2GFP-targeting sgRNAs and the thermal switch driving KRAB-dCas9. Using a NIR laser, we remotely and noninvasively maintained a focal skin temperature of tissue phantoms at either 37°C or 44°C for 30 min (Figure 2d, e). Recovery of the engineered cells 2 days following thermal treatment revealed that heated cells exhibited a significant decrease in d2GFP fluorescence compared to cells extracted from unheated phantoms (Figure 2f), demonstrating remote control of transcriptional activity in mammalian cells at distinct anatomical sites.

Here, we developed a tunable, heat-triggered platform to regulate mammalian cell transcription by remote control. Our data demonstrate that discrete pulses of heat selectively trigger the expression of dCas proteins, which then reversibly activate or suppress target genes depending on the strength and duration on the thermal inputs. Importantly, we demonstrate that transcriptional activity can be modulated by remote control in living mice. This framework establishes a noninvasive and targeted approach to harness CRISPR-based technologies for modulation of gene expression and expands the current suite of small molecule and light-based methods for dynamic control of cell function. Looking forward, incorporating multiple gene targets as well as genetic programs consisting of orthogonal commands, such as by leveraging the use of ‘dead’ sgRNAs^25^, could provide a highly multiplexed approach to remotely interrogate mammalian biology and modulate synthetic cellular phenotypes directly *in vivo*.

## MATERIALS AND METHODS

### Cell Culture

Human embryonic kidney (HEK) 293T cells were obtained from ATCC and maintained in Dulbecco’s Modified Eagle’s Medium (DMEM, Life Technologies 11995073) supplemented with 10% fetal bovine serum (FBS, Thermo Fisher 16140071), 25 mM HEPES (Thermo Fisher 15630080), and 1% penicillin/streptomycin (Life Technologies, 15140-122). All cells were cultured at 37°C in 5% CO_2_ unless otherwise noted.

### Plasmid Design and Construction

All DNA constructs (**Supplementary Figure 1**) used were delivered to cells as plasmids. Unless otherwise noted, all restriction enzymes were obtained for New England Biolabs (Ipswich, MA). All primers and sequencing verifications were obtained from Eurofins Genomics (Louisville, KY).

The hU6 promoter and the activating guide scaffold (sgRNA 2.0)^21^ (IDT PAGE Ultramer) were simultaneously inserted into the pUC57 cloning vector (GenScript SD1176) by restriction enzyme digest using EcoRI, NsiI, and BamHI. The NsiI site was then removed using the Q5 Site-directed mutagenesis Kit (New England Biolabs, E0554S). All sgRNA sequences for activation studies (**Supplementary Table 1a**) were annealed and inserted via the BbsI restriction site. Subsequently, the hU6-sgRNA-activating scaffold sequences corresponding to endogenous genes were transferred into LeGO-C with constitutive expression of blue fluorescent protein, previously inserted using the BamHI and EcoRI restriction sites. The MPH construct (Addgene #61423) containing constitutive expression of eGFP was used for all activation experiments.

sgRNA sequences for suppression studies (**Supplementary Table 1b**) were inserted via the BbsI restriction site into eSpCas9(1.1) (Addgene #71814), which contains the human U6 promoter (hU6) and a standard guide scaffold (sgRNA 1.0)^21^. Using the XhoI and ApaI restriction sites, the hU6-sgRNA-suppresion scaffold sequences were subsequently transferred into the LeGO-C lentiviral backbone (Addgene #27348) with constitutive expression of a destabilized GFP variant (d2GFP), previously inserted using the BamHI and EcoRI restriction sites.

The promoter of the HSPA6 gene (Uniprot P17066) was amplified from human genomic DNA (Clontech #636401) from −1231 bp to +119 bp relative to the transcriptional start site, as previously described^15^. The dCas9 variants (Addgene 47107, 21916) were placed under the control of the heat shock promoter via restriction enzyme cloning using AgeI and XhoI in LeGO-C. For d2GFP suppression studies, the mCherry reporter was replaced with Thy1.1 (Uniprot P01831) (**Supplementary Figure 1**).

### Validation of sgRNAs for d2GFP Suppression

One day prior to transfection, d2GFP-expressing HEK293T cells were seeded in a 6-well plate at a density of 2.5 × 10^5^ cells per well. eSpCas9(1.1) plasmids containing sgRNA sequences targeting the first 100 bps following the TSS (**Supplementary Table 1**) were transfected. Briefly, 3 μg DNA was added to 120 μL Opti-MEM medium (Life Technologies, 31985062), followed by the addition of 275 μL of Opti-MEM containing 10 μL of Lipofectamine 2000 (Invitrogen, 11668019). The mixture was incubated for 20 min at RT prior to the addition to the cells. Medium was replaced after 24 hrs with fresh DMEM (10% FBS, 1% Penicillin/Streptomycin, 25 mM HEPES). At indicated timepoints, cells were assayed for d2GFP knockout using a benchtop flow cytometer (Accuri C6 Plus, BD Biosciences). The sgRNA with most potent d2GFP knockout (Guide A3, **Supplementary Figure 6**) was used for all subsequent suppression studies.

### Viral Production and Generation of Stable Cell Lines

Plasmid DNA was purified using with either E.Z.N.A.^®^ Endo Free Plasmid Mini Kit II or Midi Kit (Omega Bio-Tek D6950-01, D6915-03) and packaged into lentiviral vectors with psPAX2 (Addgene #12260) and pMD2.G (Addgene #12259) using TransIT-LT1 transfection reagent (Mirus Bio MIR2300) and HEK293T cells. Viral supernatant was concentrated using PEG-*it* Virus Precipitation Solution (System Biosciences, Palo Alto, CA) following the manufacturer’s protocol. Cells were transduced in 10 μg/mL of protamine sulfate (Sigma P3369) with the appropriate combination of plasmids (see **Supplementary Figure 1**), depending on the application. Cells were purified via FACS (BD FACS Fusion).

### Intracellular Staining of dCas9 and Cas9

Cells stably transduced with the heat inducible portion of the circuit (*HSPA6-KRAB dCas9-IRES-mCherry SFFV Thy 1.1* or *HSPA6-Cas9-IRES-mCherry SFFV Thy 1.1*, **Supplementary Figure 1**) were heated in a thermal cycler (42°C, 30 min, unless otherwise noted) at a density of 10^6^ cells/mL and immediately transferred to a 6-well plate at 3×10^5^ cells/well in triplicates. At the indicated timepoints, triplicate wells were trypsinized and washed twice with 1× PBS(-/-). Cells were fixed for 60 min at RT with the Foxp3 Fixation/Permeabilization Buffer (Invitrogen 00552300), washed, and blocked with 3.5% BSA solution for 45 min. Cells were incubated for 30 min at RT with primary antibodies (Biolegend 844301, anti-Cas9 Clone 7A9) diluted at 1:100, and subsequently washed twice prior to incubation in the dark at RT with suitably matched secondary antibodies (Biolegend 406605, FITC anti-mouse IgG1 Clone RMG1-1) for an additional 45 min. Cells were washed again before analysis by flow cytometry (Accuri C6 Plus, BD Biosciences.)

### In Vitro Heating Assays for d2GFP Suppression

Cells were heated in a thermal cycler (42°C, 30 min) at a density of 10^6^ cells/mL, immediately transferred to a 24-well plate at 5×10^5^ cells/well in triplicates, and incubated at 37 °C and 5% CO_2_. At the indicated timepoints, cells were trypsinized and analyzed by flow cytometry (**Supplementary Figure 7**, BD Accuri C6). Cells were passaged when they reached approximately 70% confluence. For reheating assays, cells were trypsinized and heated once again at the indicated timepoints.

### In vivo suppression of d2GFP

All animal studies were approved by the Institutional Animal Care and Use Committee at Georgia Institute of Technology. Gold nanorods (AuNRs) were synthesized as previously described^15^. 0.5 μg AuNRs and 2.5×10^5^ cells were mixed into 100 μL Matrigel^®^ (Corning 354248, final concentration 8 mg/mL) and injected subcutaneously into Nude mice (002019 NU/J, The Jackson Laboratory). Three days following implantation, implant sites were heated using an 808 nm laser (Coherent) at a power density of ∼9.5 A/cm^2^. The surface temperature of heated implants was maintained at 44 ± 1°C for 30 minutes and continually monitored using a thermal camera (FLIR 450sc). Two days post-heating, Matrigel implants were excised and incubated with 350 μL of cell recovery solution (Liberase DL [0.5 mg/mL, Sigma 5401160001] and DNAse I [0.1 mg/mL, 10104159001] in Opti-MEM [Life Technologies, 31985062]) for 30 min at 37°C. Recovered cells were analyzed by flow cytometry (BD Accuri C6).

### qRT-PCR analysis of transcriptional activation

One day prior to transfection, HEK293T cells stably expressing plasmids in **Supplementary Figure 1a** were seeded in a 6-well plate at a density of 5.0 × 10^5^ cells per well. sgRNA-coding plasmids targeting the ~200 bp window upstream of the TSS (**Supplementary Table 1**) were pooled together for each gene and subsequently transfected at a 1:1:1 mass ratio with plasmids described in **Supplementary Figure 1a**. Briefly, 3 μg total DNA was added to 120 μL Opti-MEM medium, followed by the addition of 275 μL of Opti-MEM containing 10 μL of Lipofectamine 2000. The mixture was incubated for 20 min at RT prior to the addition to the cells. 24 hrs following transfection, cells were heated for 30 minutes at the indicated temperatures. mRNA was collected 48 and 72 hrs post heating and purified using the RNeasy Plus Mini Kit (Qiagen 74134). cDNA was synthesized from 0.4 μg of total cellular RNA using RT^2^ First Strand Kit (Qiagen 330404). TaqMan qPCR probes (Thermo Fisher Scientific, **Supplementary Table 3**) and Fast Advanced Master Mix (Thermo Fisher Scientific, 4444556) were used in 10 μL reactions in quadruplicates in a 384-well format. Relative levels of cDNA were detected using QuantStudio^TM^ 6 Flex Real-Time PCR System (Applied Biosciences). Raw data was normalized to GAPDH levels and untreated (37°C) controls using the ΔΔCt method. Analyzed data are reported as mean ± s.d.

### Statistical analysis

All results are presented as mean and error bars show standard deviation. Statistical analysis was performed using statistical software (GraphPad Prism 6; GraphPad Software). \**P* < 0.05, ** *P* < 0.01, *** *P* < 0.001.

## ACKNOWLEDGEMENTS

We thank Dr. J.E. Dahlman (Georgia Tech), Dr. D.R. Meyers (Emory), and C.D. Sago (Georgia Tech) for helpful insights. This work was funded by the NIH Director’s New Innovator Award (DP2HD091793), the National Center for Advancing Translational Sciences (UL1TR000454), the Shurl and Kay Curci Foundation, and was partially performed at the Georgia Tech Institute for Nanotechnology, a member of the National Nanotechnology Coordinated Infrastructure which is supported by the National Science Foundation (Grant ECCS-1542174). L.G. is supported by the Alfred P. Sloan Foundation, the National Institutes of Health GT BioMAT Training Grant under Award Number 5T32EB006343 and the National Science Foundation Graduate Research Fellowship under Grant No. DGE-1451512. G.A.K. holds a Career Award at the Scientific Interface from the Burroughs Wellcome Fund. This content is solely the responsibility of the authors and does not necessarily represent the official views of the National Institutes of Health.

## AUTHOR CONTRIBUTIONS

L.G. and G.A.K. conceived of the idea. L.G. and G.A.K. designed the experiments and interpreted results. L.G., E.V.P., H.L., and J.P.M performed the experiments. L.G. and G.A.K. wrote the manuscript.

## DATA AVAILABILITY

The data that support the findings of this study are available from the authors upon reasonable request.

