## Supplementary Information for "Heat-triggered remote control of CRISPR-dCas9 for tunable transcriptional modulation"

Address: Marcus Nanotechnology Building, 345 Ferst Drive, Atlanta, GA 30332, USA

### SUPPLEMENTARY RESULTS

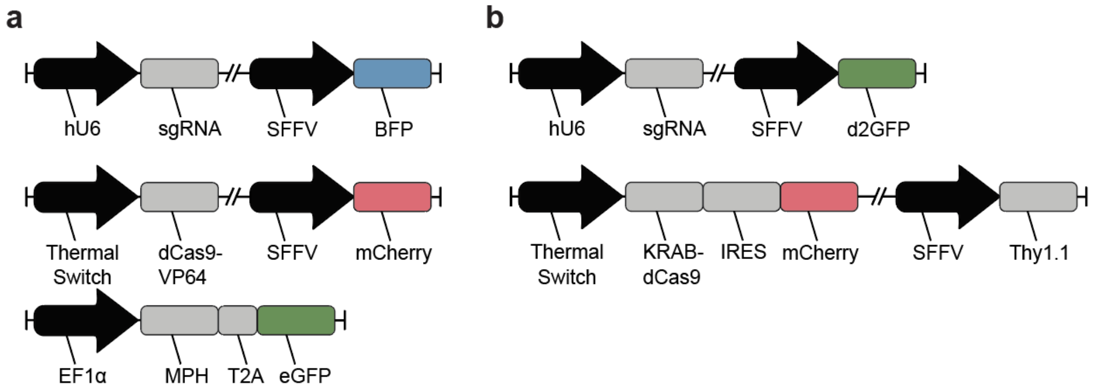


**Supplementary Figure 1 | Genetic circuits for thermal control of transcription.**

DNA constructs for thermal control of (a) transcriptional activation of endogenous genes and (b) suppression of d2GFP. Guide RNAs (sgRNAs) are constitutively expressed via the human U6 (hU6) promoter. The thermal switch controls the expression of dCas9 variants (dCas9-VP64, KRAB-dCas9). For activation studies, a third plasmid coding for the MS2-P65-HSF1 (MPH) transcriptional activation complex is included, which is reported to regulate efficient target upregulation by binding to MS2 loops included within the sgRNA scaffold[1].

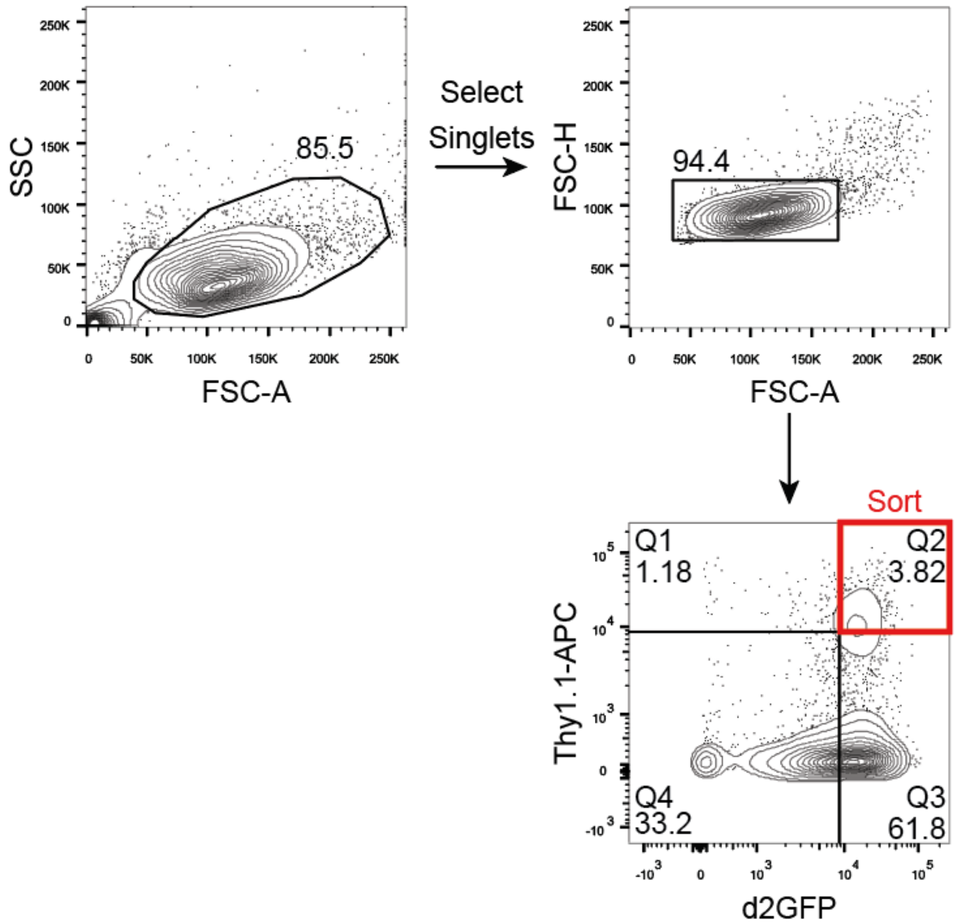


**Supplementary Figure 2 | Gating strategy for selecting HEK293T cells for d2GFP suppression .** Gating strategy used for selecting cells that are double positive for the thermal switch (reporter: Thy1.1) and sgRNAs (reporter: d2GFP).

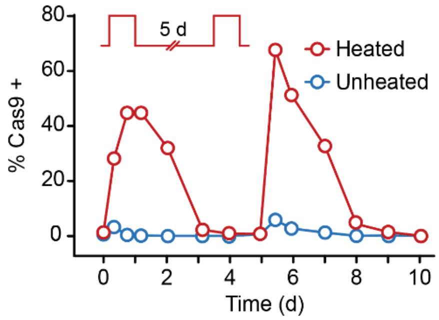


**Supplementary Figure 3 | Short heat pulses enable transient control of catalytically-active Cas9 expression.** Kinetic trace of Cas9 expression in HEK293Ts triggered by the HSPA6 thermal switch. Cells were heated to 42°C for 30 min at t=0 d and t= 5 d.

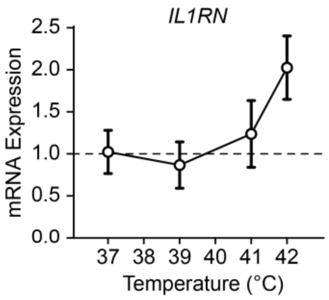


**Supplementary Figure 4 | Modulation of endogenous gene activation by varying temperature.** Endogenous gene activation as assessed by qRT-PCR 72 hrs after heating at the indicated temperatures (*n* = 3-4, mean ± s.d.)

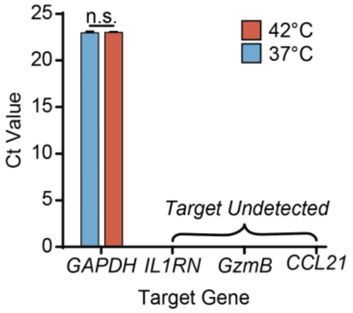


**Supplementary Figure 5 | Heating wild-type HEK293Ts does not activate chosen endogenous genes.** *IL1RN*, *GzmB* and *CCL21* transcripts are undetected in wildtype HEK293Ts heated at 37°C and 42°C for 30 min, assayed 72 hrs post heating (*n* = 4, mean ± s.d., unpaired t-test, *P<0.05)

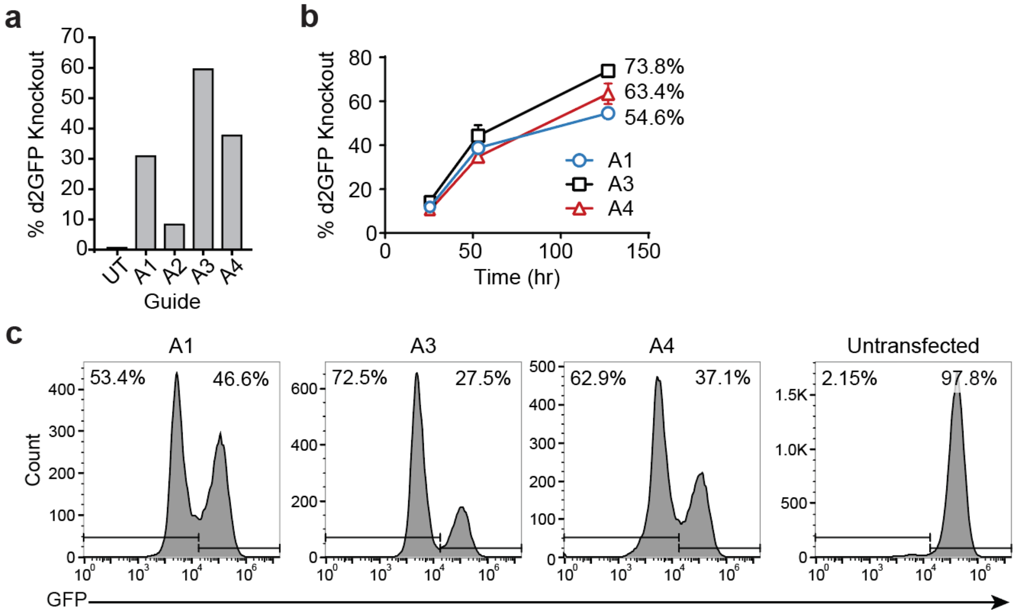


**Supplementary Figure 6 | Scouting sgRNAs for d2GFP Suppression.** eSpCas9(1.1) plasmids containing sgRNA sequences targeting the first 100 bps following the translational start site (Supplementary Table 2) were transfected into d2GFP-expressing HEK293T cells. (a) d2GFP knockout was measured by flow cytometry 9 days post transfection. (b) The kinetics of d2GFP knockout were measured on the top performing guides (*n*=*3*, mean ± s.d.). (c) Representative flow plots of d2GFP expression 127 hrs post transfection.

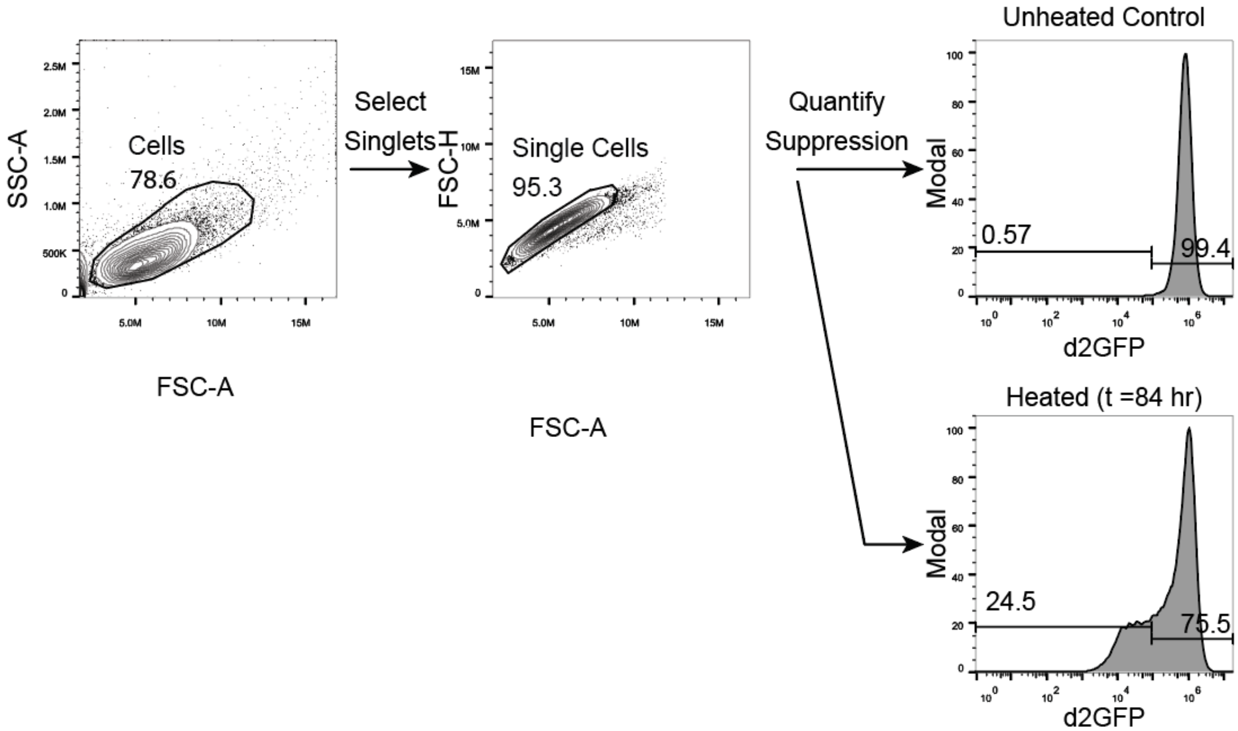


**Supplementary Figure 7 | Gating strategy for quantifying d2GFP suppression.**

Representative gating strategy used for quantifying d2GFP suppression. Representative histograms shown for unheated cells (top right) and heated cells (bottom right).

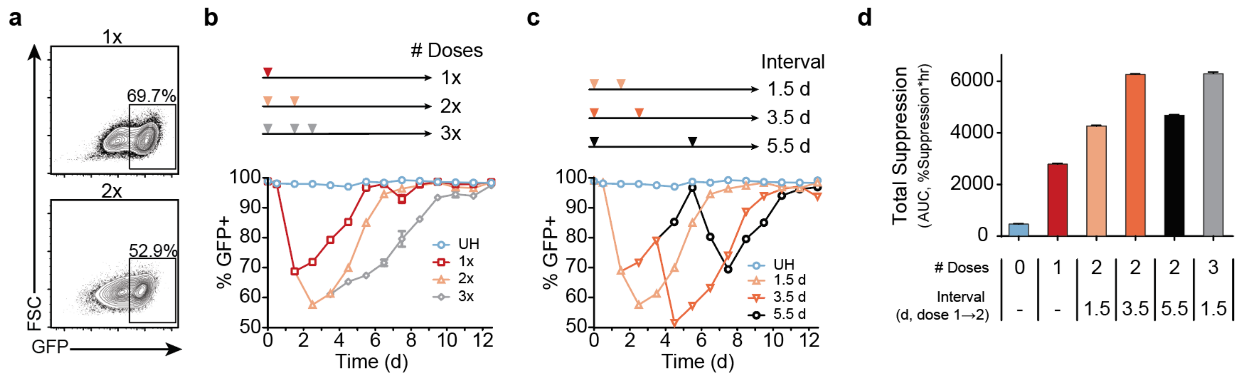


**Supplementary Figure 8 | Heating frequency and interval impact d2GFP suppression kinetics.** Expression of d2GFP was assessed by flow cytometry in HEK293T transduced with thermal control of KRAB-dCas9 driven transcriptional suppression (Supplementary Figure 1b). (a) Representative flow plots of maximal d2GFP suppression following 1 (top) and 2 doses (bottom) of heat. (b) Kinetic trace of d2GFP expression following 1, 2, or 3 heat doses (c) or following 2 heat doses delivered at different intervals (1.5, 3.5, or 5.5 d). (d) Total suppression measured for all conditions. All heat treatments were administered for 30 min at 42°C (*n=3*, mean ± s.d.).

**Supplementary Table 1 |** sgRNA sequences for transcriptional activation.

| Target Gene | Strand | Guide # | Sequence (5'-3') |
|---|---|---|---|
| CCL21 | Top | 1 | CACCGGGTAGCTGGGAATAGAAGGA |
| | | 2 | CACCGGAGGGGAAGGGTATGGATCC |
| | | 3 | CACCGGAGACAGTCATGGTGTTCCA |
| | | 4 | CACCGGACATAAAATTTGGCAGCTG |
| | | 5 | CACCGGCGTAGTGAGGAGACAGTCA |
| | Bottom | 1 | AAACTCCTTCTATTCCCAGCTACCC |
| | | 2 | AAACGGATCCATACCCTTCCCCTCC |
| | | 3 | AAACTGGAACACCATGACTGTCTCC |
| | | 4 | AAACCAGCTGCCAAATTTTATGTCC |
| | | 5 | AAACTGACTGTCTCCTCACTACGCC |
| GZMB | Top | 1 | CACCGGGCACCCAGAGGACGTCATC |
| | | 2 | CACCGGAGAGGACGTCATCAGGCAG |
| | | 3 | CACCGGTCAGCTGTGGGTGATGATG |
| | | 4 | CACCGGACTCTGAGTCATCAGCTGT |
| | | 5 | CACCGGCTGCTCTGGGCTGAATAGG |
| | Bottom | 1 | AAACGATGACGTCCTCTGGGTGCCC |
| | | 2 | AAACCTGCCTGATGACGTCCTCTCC |
| | | 3 | AAACCATCATCACCCACAGCTGACC |
| | | 4 | AAACACAGCTGATGACTCAGAGTCC |
| | | 5 | AAACCCTATTCAGCCCAGAGCAGCC |
| IL1RN | Top | 1 | CACCGGTGTACTCTCTGAGGTGCTC |
| | | 2 | CACCGGACGCAGATAAGAACCAGTT |
| | | 3 | CACCGGCATCAAGTCAGCCATCAGC |
| | | 4 | CACCGGGAGTCACCCTCCTGGAAAC |
| | Bottom | 1 | AAACGAGCACCTCAGAGAGTACACC |
| | | 2 | AAACAACTGGTTCTTATCTGCGTCC |
| | | 3 | AAACGCTGATGGCTGACTTGATGCC |
| | | 4 | AAACGTTTCCAGGAGGGTGACTCCC |

**Supplementary Table 2 |** sgRNA sequences for d2GFP suppression.

| Target Gene | Strand | Guide ID | Sequence (5'-3') |
|---|---|---|---|
| d2GFP | Top | A1 | CACCGCGAGGAGCTGTTCACCGGGG |
| | | A2 | CACCGCACCGGGGTGGTGCCCATCC |
| | | A3 | CACCGGACCAGGATGGGCACCACCC |
| | | A4 | CACCGAAGGGCGAGGAGCTGTTCAC |
| | Bottom | A1 | AAACCCCCGGTGAACAGCTCCTCGC |
| | | A2 | AAACGGATGGGCACCACCCCGGTGC |
| | | A3 | AAACGGGTGGTGCCCATCCTGGTCC |
| | | A4 | AAACGTGAACAGCTCCTCGCCCTTC |

**Supplementary Table 3 |** TaqMan® probes used for qRT-PCR assays.

| Target Gene | Taqman ® Gene Expression Assay ID |
|---|---|
| IL1RN | Hs00893626_m1 |
| GZMB | Hs00188051_m1 |
| CCL21 | Hs00171076_m1 |
| GAPDH | Hs03929097_g1 |
